# Genetic and morphological variation is associated with differences in cold-water tolerance and geographic expansion among invasive snails (*Melanoides tuberculata*) in central Texas

**DOI:** 10.1101/2019.12.20.884866

**Authors:** Stephen Harding, David Rodriguez, Jacob Jackson, David Huffman

**Author notes:** Corresponding author: David Rodriguez, Department of Biology, Texas State University, 601 University Drive, San Marcos, TX 78666, USA, E-mail address.

## Abstract

*Melanoides tuberculata* (Thiaridae) is an old-world freshwater snail that is and now circumtropical. After being introduced in the 1960s via the aquarium trade, populations of *M. tuberculata* are thriving in spring systems of Texas (USA). Field surveys and experimental investigations of temperature tolerance suggest *M. tuberculata* is stenothermal, and thus range expansions outside of aquatic habitats with water temperatures between 18° and 32°C should be unlikely. However, in 2012 snails were detected in natural aquatic habitats with seasonal temperatures below the experimentally determined lethal thermal minimum. To test whether genetic and phenotypic variation might be associated with cold-water tolerance and range expansion, we sequenced the 16S ribosomal rRNA gene and measured qualitative conch morphology of 170 snails collected at 26 sites in three central Texas rivers. We also conducted phylogenetic analyses of *M. tuberculata* collected globally and in Texas to determine potential source populations and estimate the number of invasion events. Our results show snails detected in variable temperature habitats are genetically divergent and morphologically distinct from snails collected in habitats with stable temperatures. These data are consistent with at least three introduction events into Texas of *M. tuberculata sensu lato* that are characterized by distinct habitat preferences, physiological tolerances, and/or adaptive behaviors.

## Introduction

As invaders, mollusks can have costly and dramatic effects on invaded ecosystems. For example, repair and maintenance costs of industrial pipes and drains as a consequence of zebra mussel *(Dreissena polymorpha)* and Asian clam *(Corbicula fluminea)* biofouling are estimated to cost local municipalities hundreds of millions in the US (U.S. Congress Office of Technology Assessment 1993; Nelson 2019). Invasive mollusks can compete with native species (Riley et al. 2008), have been implicated in water quality degradation (Effler et al. 1996), and have contributed to extirpation of native species (Ricciardi et al. 1998). Additionally, some gastropods serve as intermediate hosts in the life cycles of parasites that in turn pose health risks to humans and animals (Keawjam et al. 1993; Pokora 2001; Pinto and De Melo 2011). Therefore, preventing the spread of invasive mollusks is economically and ecologically important, and understanding the life history of invasive taxa in non-native habitats is paramount to mitigating their expansion.

In general, successful invaders are generalists, strong competitors, have high fecundity rates, and can resist wide fluctuations in environmental conditions (Sakai et al. 2001). *Melanoides tuberculata* is an aquatic snail native to Africa, Asia, and the Middle East but is now circumtropical. These snails can reproduce sexually or via parthenogenesis; although, males are typically rare or completely absent in some populations (Livshits and Fishelson 1983; Heller and Farstey 1990). They also have high fecundity rates (Freitas et al. 1987; Work and Mills 2013) and can tolerate broad environmental conditions with the exception of sensitivity to cold water (Mitchell and Brandt 2005; Mitchell et al. 2007). They also serve as an intermediate host for more species of trematode parasites than any other aquatic snail reported, including 11 in Asia that infect humans (Pinto and De Melo 2011); consequently, the presence of *M. tuberculata* in invaded systems is a concern for wildlife conservation and human health.

Thiarids entered North America as early as 1935 via the aquarium trade (Murray and Wopschall 1965). A specific timeline for the introduction of *M. tuberculata* is unknown; however, evidence suggests they were introduced in the same manner (Roessler et al. 1977; Neck 1985). *Melanoides tuberculata* was first detected in central Texas rivers in the 1960s and 70s (San Antonio River, Bexar County, 1965; Comal River, Comal County, 1965; San Marcos River, Hays County, 1979) (Murray and Wopschall 1965; Lindholm 1979) and then later in multiple spring-fed systems throughout the continental USA (Murray 1975; Rader et al. 2003; Wingard et al. 2008; Karatayev et al. 2009; Daniel et al. 2019). Together these studies suggest that central Texas was potentially the epicenter of invasion into North America for *M. tuberculata.*

Field surveys in the Bonneville Basin, Utah (USA) detected *M. tuberculata* at multiple locations, except in springs with water temperatures below 18°C (Rader et al. 2003). Their observations were consistent with other field surveys, which suggest suitable habitat is limited to waters with little to no flow velocity and stable temperatures between 18°C and 30°C (Murray 1971; Neck 1985). Controlled experiments investigating temperature tolerance presented evidence that survival is unlikely when *M. tuberculata* is exposed to water temperatures below 18°C or above 32°C for more than 48 hours (Mitchell and Brandt 2005). Thus, it would appear that any further range expansions of *M. tuberculata* into temperate aquatic habitats with seasonal temperatures exceeding these limits would be unlikely owing to their expected thermal tolerances.

In 2000 and 2001, three searches for *M. tuberculata* in the Guadalupe River (GR) (New Braunfels, Texas), near its confluence with the Comal River (CR), yielded no live snails (Fleming et al. 2011). These survey results were consistent with expectations, because seasonal water temperatures fell below 18°C in this stretch of river. However, a subsequent search in 2009 detected *M. tuberculata* thriving in the GR up to 2 km downstream from the confluence (Huffman pers. comm.). In 2010, live snails measuring > 50 mm were found in the GR approximately 15 km downstream from the CR (Huffman pers. comm.; Figure 1). The size of these snails suggests they were several years old (Pointier et al. 1993) and represent an invasive range expansion. Until recently, the known range of *M. tuberculata* in the San Marcos River (SMR) was within three river km downstream from the headwaters (Figure 1), and it has never been reported outside of the spring influenced reach (Huffman pers. comm.). In 2014, snails morphologically identified as *M. tuberculata* were thriving in the lower SMR approximately 54 river km downstream from the spring-fed headwaters (Figure 1). Expansion of *M. tuberculata* into these areas is unusual, because these habitats have minimum seasonal temperatures as low as 10°C, which is well below the experimentally determined minimum temperature of 18°C for this species (Figure 1; Table 1).

**Figure 1.**
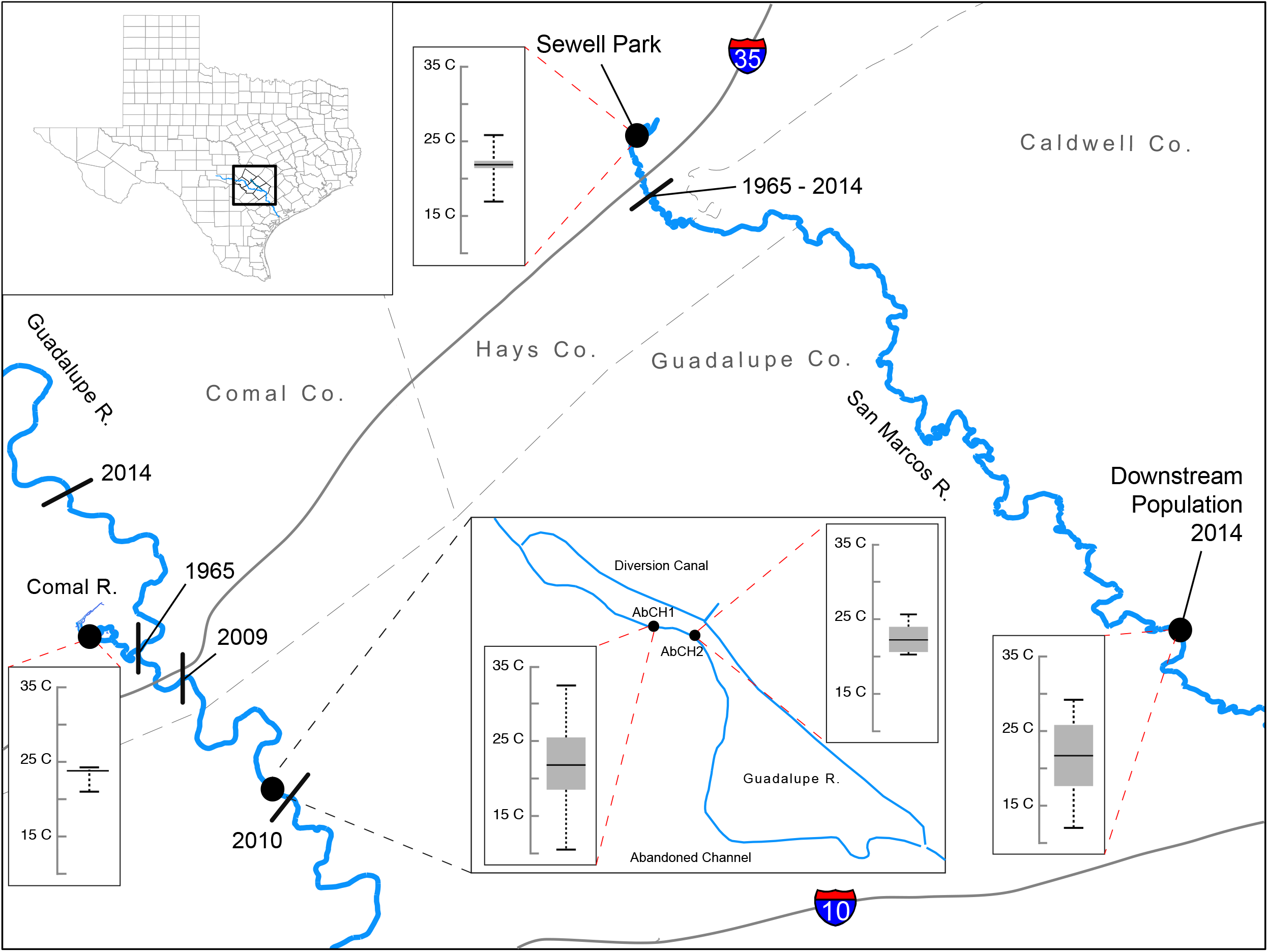
Known distributions and range expansion of *Melanoides tuberculata* in central Texas (black bars represent the furthest range extent during a survey year. Box plots show mean water temperature (°C) data (black bar) collected at different sites in the Comal (Landa Lake), Guadalupe, Upper San Marcos (near Spring Lake), and Lower San Marcos rivers, with minimum and maximum temperatures (whiskers) and upper and lower quartiles (grey box).

**Table 1.** Summarized measures of central tendency and dispersion for water temperature data collected at each site in central Texas (SMR HW = San Marcos River headwaters, SMR 13 = San Marcos River site 13, ULL = Upper Landa Lake, LLL = Lower Landa Lake, AbCH1 = Abandoned Channel site 1 Guadalupe River, and AbCH2 = Abandoned Channel site 2 Guadalupe River). Temperature was measured in degrees Celsius (sd = standard deviation).

| Site | Mean | Median | Minimum | Maximum | sd |
| --- | --- | --- | --- | --- | --- |
| SMR HW | 21.914 | 21.868 | 16.924 | 25.833 | 0.783 |
| SMR 13 | 21.649 | 21.7 | 12.025 | 29.19 | 4.628 |
| ULL | 23.593 | 23.678 | 19.969 | 25.028 | 0.318 |
| LLL | 23.747 | 23.803 | 21.007 | 24.264 | 0.192 |
| AbCH1 | 21.936 | 21.795 | 10.516 | 32.407 | 4.806 |
| AbCH2 | 22.425 | 22.226 | 20.15 | 25.525 | 1.733 |

Global populations of *M. tuberculata* are characterized by a broad range of shell morphologies and high genetic variation among clonal lineages, but comparatively low variation within clonal lineages (Livshits et al. 1984; Samadi et al. 1998; Samadi et al. 1999; Samadi et al. 2000; Facon et al. 2003; Genner et al. 2004; Sørensen et al. 2005; Van Bocxlaer et al. 2015). Previous phylogenetic treatments present evidence for polyphyly within *M. tuberculata sensu lato* (Facon et al. 2003; Genner et al. 2004; Sørensen et al. 2005; Van Bocxlaer et al. 2015), which suggests strong lineage divergence within the species. Also, conferred advantageous physiological limits have been observed in offspring of hybridized lineages of *M. tuberculata* that were not possessed by the parental snails (Facon et al. 2005). Comprehensively, these reports serve as compelling evidence that genetic structure within and among the *M. tuberculata sensu lato* (e.g., genotype and/or haplotype) can be associated with differences in life history and/or physiological limits.

Lastly, field surveys in Israel found *M. tuberculata* survived during colder months by hibernating in soft sediments (Livshits and Fishelson 1983), which provides some evidence of adaptive behavioral differences within the *M. tuberculata* species group. However, the association between phylogeny and this behavior is still unknown. Accordingly, measuring, documenting, and reporting intra-specific and inter-population level variation across the landscape is important, because it provides evolutionary context when differences in adaptive potential and behaviors are exhibited by genetic variants. Thus, expansion of *M. tuberculata sensu lato* into Texas waters that seasonally reach temperatures below experimentally determined lethal thermal limits might be explained by the introduction of different lineages already adapted to colder temperatures in their native environments.

To investigate the introduction and expansion of *M. tuberculata* in central Texas, we aim to measure variation among contemporary populations via combined genetic and morphological analyses. We also aim to delimit the distribution of unique haplotypes and morphotypes while providing geographic ranges and habitat associations for each. Lastly, we aim to estimate the evolutionary history of haplotypes detected and attempt to infer their geographic origin based on global reference data.

## Methods

### Water temperature data collection

We used HOBO data loggers to collect temperature data from two sites in the San Marcos River, one site in the Comal River, and two sites in the Guadalupe River (Figure 1). In some cases, the onboard logger memory reached capacity before we extracted the data, which resulted in some gaps at various intervals. These gaps did not affect our temperature estimates. We plotted calculated mean temperature and standard deviation as well as raw temperature data using *ggplot2* package (Wickham 2016) in R (Figure S1). We calculated and summarized measures of central tendency and dispersion (Table 1). We also plotted the minimum, maximum, and mean temperatures, as well as the upper and lower quartiles using the *boxplot* function in R (Figure 1). In March 2013, the logger at AbCH2 was lost following a flood event; therefore, we only plotted the data before this event (Figure S1).

### Sampling

We sampled a total of 26 publicly accessible locations along three central Texas rivers, which included eight sites on the CR, seven sites on the upper SMR, nine sites on the lower SMR (to Luling, TX), and two on the GR (Figure 1; Table 2). Sampling efforts in the GR were restricted by increased amounts of rainfall in central Texas from 2015 to mid-2016. Flooding and high flow velocities restricted access to many of the areas in the GR that previously held populations of *M. tuberculata.* Subsequent searches after the river returned to base flows yielded no live snails at several localities, thus those sites were omitted from this study. The two sites within the GR included in this study were sampled in early 2017.

**Table 2.**
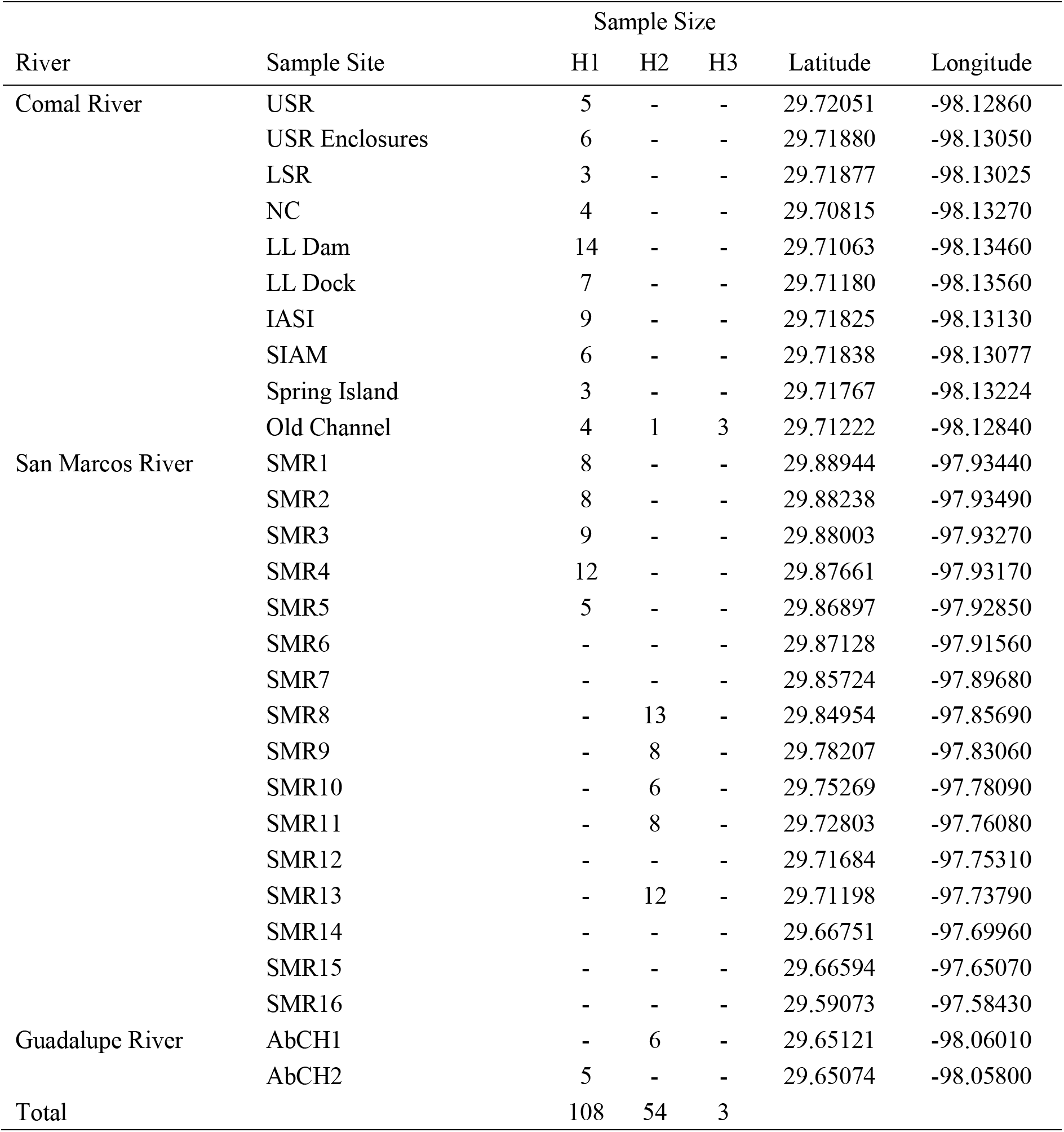
Rivers sampled for *Melanoides tuberculata*, sites within each river, sample size for each 16S haplotype, and GPS coordinates for each site. Codes for each sampled site: USR = Upper Spring Run, LSR = Lower Spring Run, NC = New Channel, LL DAM = Landa Lake near the dam, LL Dock = Landa Lake near the paddle boat dock, IASI = Island Above Spring Island, SIAM = Spring Island Above Mulberry Street, SMR = San Marcos River, AbCh = Abandoned Channel (see Figure 2).

We searched for snails in shallow, slack water riparian habitats using dip nets and used snorkeling and scuba diving to search deeper habitats. Sampling intensity ranged from two to four people searching for one hour at each site. Due to variable snail densities among sites and storage limitations, we chose to retain up to 20 live snails from each site. If we collected more than 20 snails at a site, then we subset the collection by choosing representatives of the different phenotypes from the total snails collected at that site. Our goal was not to measure density among sites, but rather to estimate the breadth of phenotypic and genetic diversity across rivers. We promptly transported all snails to Texas State University where they were housed in flowthrough containers submerged in flowing artesian water (maximum of seven days) until we performed morphometric and genetic analyses.

### Genetic Analyses

We used approximately 25 mg of tissue collected from either the head or foot of the snail. We used two separate extraction kits during the study: the Gentra PureGene Tissue Kit (Qiagen) and the GeneJET Genomic DNA Purification Kit (Thermo Fisher Scientific), following each respective manufacturer’s protocols. When using the GeneJET Kit, we followed the Mammalian Tissue and Rodent Tail Genomic DNA Purification Protocol. We ensured the presence of high molecular weight DNA for each extraction using electrophoresis in a 1% agarose gel and 0.5X TBE buffer solution.

Polymerase Chain Reactions (PCR) were performed using specific *M. tuberculata* mitochondrial DNA primers targeting the 16S rRNA gene; namely, 16SF (5’-TAGCATGAATGGTCTGACGAAAGC-3’) and 16SR (5’-AAGGAGATTATGCTGTTATCCC-3’) (Facon et al. 2003). All amplifications were performed at 25 μL volumes containing 12.5 μL 2X DreamTaq PCR Master Mix (Thermo Fisher Scientific), 1 μM forward primer, 1 μM reverse primer, 0.10-20 ng/μL of genomic DNA, and nuclease free H_2_O to volume. We used a PCR thermal profile similar to the one used in Facon et al. (2003), which began with initial denaturation at 92 °C for 1 min.; then 40 cycles of 92 °C for 20 s, 48 °C for 40 s, and 68 °C for 1 min; and a final elongation at 68° for 7 mins. All reactions were diagnosed for successful amplification using a 2% agarose gel and 0.5X TBE buffer solution. We cleaned up PCR amplicons using ExoSAP-IT^®^ (Affymetrix) and cycle sequenced both DNA strands using BigDye^®^ Terminator v3.1 (Applied Biosystems Inc.). Cycle sequencing products were purified using Sephadex™ G-50 Superfine (GE Healthcare) columns (2.6 g/ 40 mL of nuclease-free water). Purified cycle sequencing products were dehydrated overnight and then incubated in 10-15 μL formamide at 95.0° for 3 mins and then sequenced on an Applied Biosystems 3500 Genetic Analyzer.

To perform sequence editing, alignment, and phylogenetic analyses, we used Geneious Pro v5.5.9 (Biomatters, Ltd.). To visualize the mutation steps between the haplotypes collected in Texas, we constructed a median-joining haplotype network using PopART (Bandelt et al. 1999). We included global 16S rRNA sequence data for *Melanoides* sp. and other thiarids accessioned into GenBank to determine the phylogenetic position of Texas snails (Table 3). Sequences downloaded from GenBank were associated with a morph code reported from their respective publications (Table 3). We assembled and trimmed sequences down to the shortest fragment in the assembly (250 bp) to generate an alignment with no missing data and performed a final alignment using the ClustalW algorithm with default values (Thompson et al. 1994; Facon et al. 2003). To determine the model of sequence evolution that best fit the data, we applied AIC in MODELTEST using the PAUP* Geneious plugin. We used Bayesian inference (MrBayes 3.2.6 plug-in; Huelsenbeck and Ronquist 2001) to infer phylogenetic relationships among globally collected *M. tuberculata* and to determine the points of origin for invasive populations in Texas. We used the best fit model and ran four heated chains for 2,000,000 iterations while subsampling every 500 and discarding a burn-in of 100,000 iterations. Priors were set to default values. Partial sequences from *Lavigeria grandis* (AY456594) and *Cleopatra bulimoides* (AY791935) represented outgroup taxa.

**Table 3.**
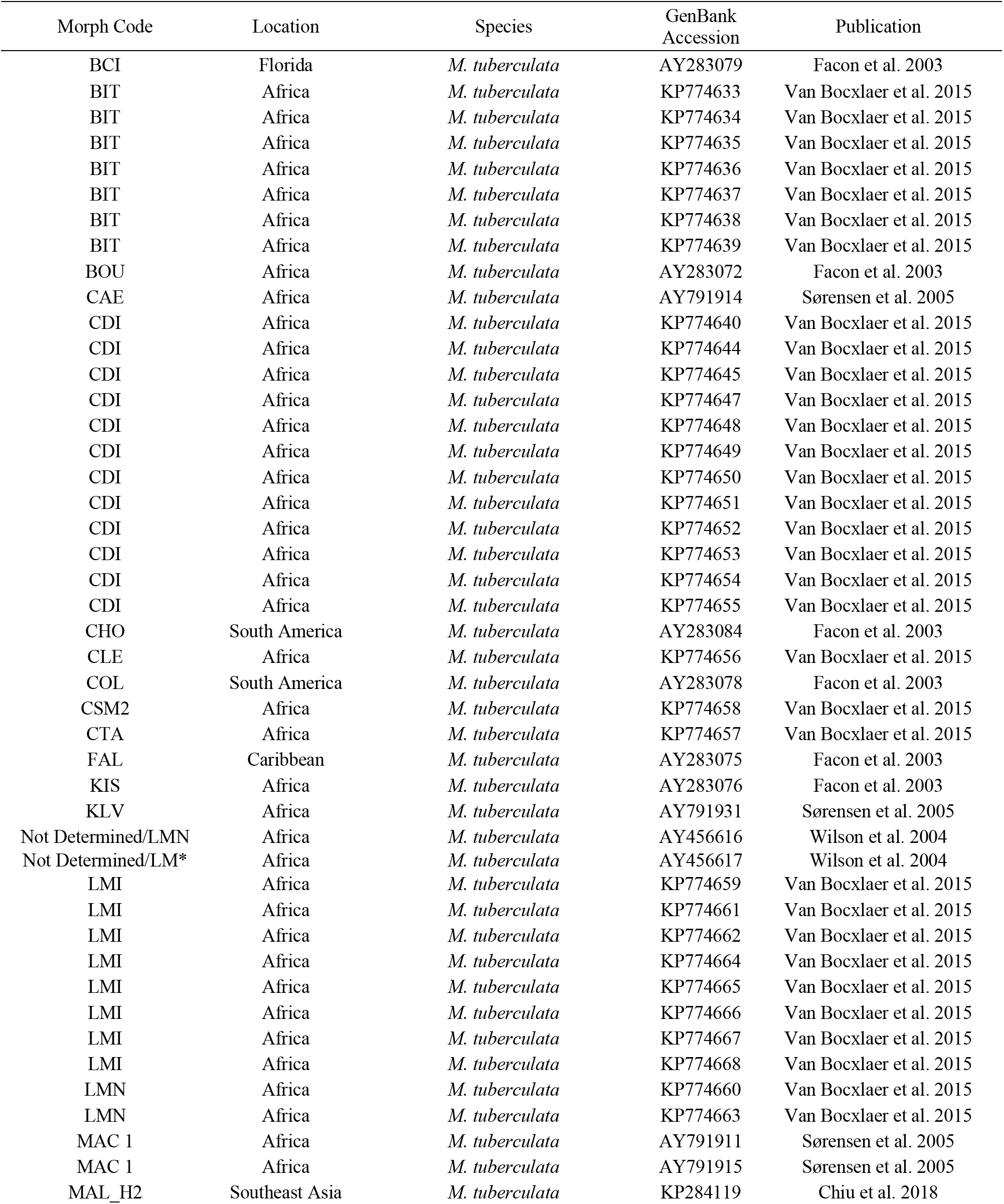
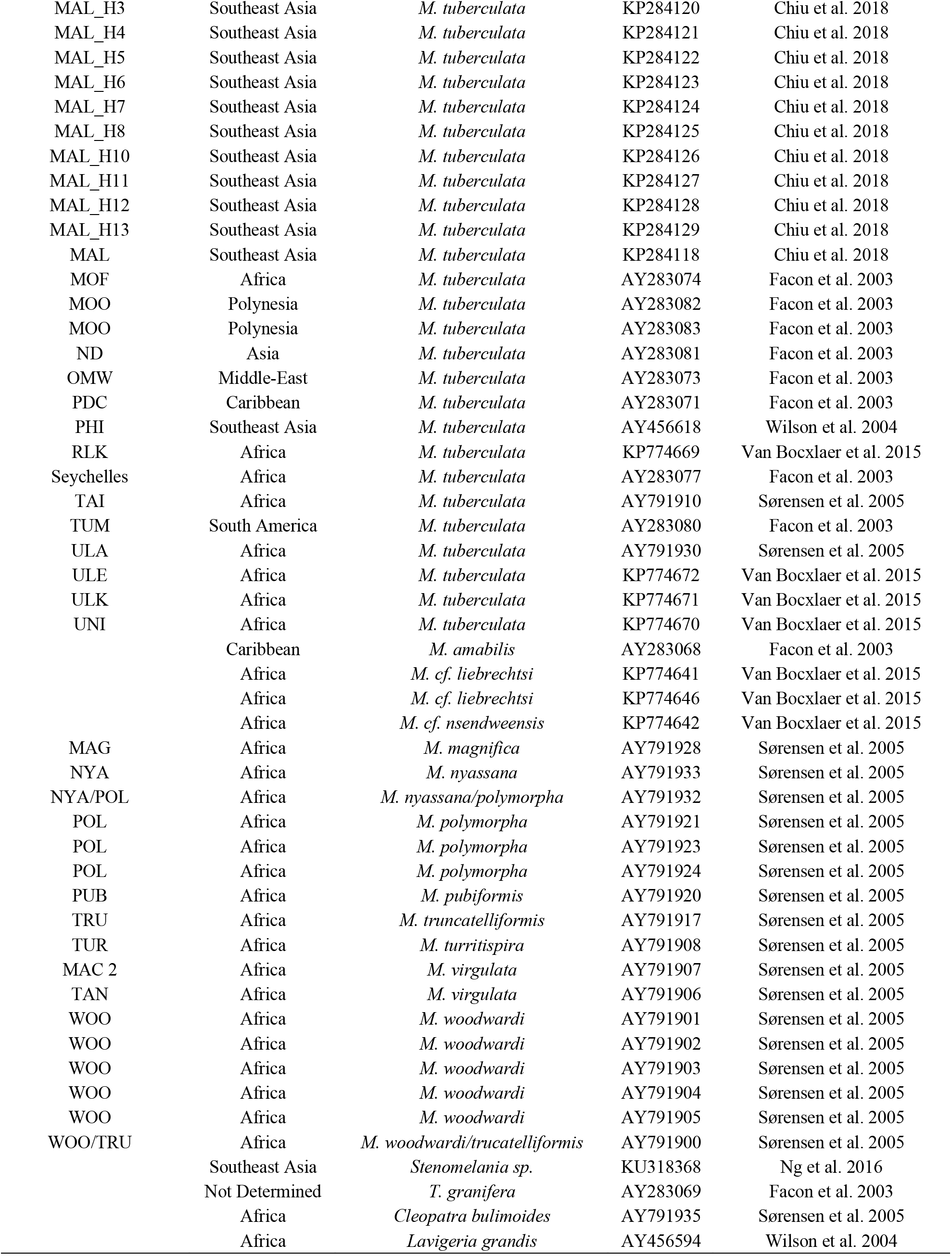
List of *Melanoides* snail morphs used in phylogenetic analyses, where they were collected, species identification, GenBank accession number, and the publication where the data were sourced.

| Morph Code | Location | Species | GenBank Accession | Publication |
| --- | --- | --- | --- | --- |
| BCI | Florida | *M. tuberculata* | AY283079 | Facon et al. 2003 |
| BIT | Africa | *M. tuberculata* | KP774633 | Van Bocxlaer et al. 2015 |
| BIT | Africa | *M. tuberculata* | KP774634 | Van Bocxlaer et al. 2015 |
| BIT | Africa | *M. tuberculata* | KP774635 | Van Bocxlaer et al. 2015 |
| BIT | Africa | *M. tuberculata* | KP774636 | Van Bocxlaer et al. 2015 |
| BIT | Africa | *M. tuberculata* | KP774637 | Van Bocxlaer et al. 2015 |
| BIT | Africa | *M. tuberculata* | KP774638 | Van Bocxlaer et al. 2015 |
| BIT | Africa | *M. tuberculata* | KP774639 | Van Bocxlaer et al. 2015 |
| BOU | Africa | *M. tuberculata* | AY283072 | Facon et al. 2003 |
| CAE | Africa | *M. tuberculata* | AY791914 | Sørensen et al. 2005 |
| CDI | Africa | *M. tuberculata* | KP774640 | Van Bocxlaer et al. 2015 |
| CDI | Africa | *M. tuberculata* | KP774644 | Van Bocxlaer et al. 2015 |
| CDI | Africa | *M. tuberculata* | KP774645 | Van Bocxlaer et al. 2015 |
| CDI | Africa | *M. tuberculata* | KP774647 | Van Bocxlaer et al. 2015 |
| CDI | Africa | *M. tuberculata* | KP774648 | Van Bocxlaer et al. 2015 |
| CDI | Africa | *M. tuberculata* | KP774649 | Van Bocxlaer et al. 2015 |
| CDI | Africa | *M. tuberculata* | KP774650 | Van Bocxlaer et al. 2015 |
| CDI | Africa | *M. tuberculata* | KP774651 | Van Bocxlaer et al. 2015 |
| CDI | Africa | *M. tuberculata* | KP774652 | Van Bocxlaer et al. 2015 |
| CDI | Africa | *M. tuberculata* | KP774653 | Van Bocxlaer et al. 2015 |
| CDI | Africa | *M. tuberculata* | KP774654 | Van Bocxlaer et al. 2015 |
| CDI | Africa | *M. tuberculata* | KP774655 | Van Bocxlaer et al. 2015 |
| CHO | South America | *M. tuberculata* | AY283084 | Facon et al. 2003 |
| CLE | Africa | *M. tuberculata* | KP774656 | Van Bocxlaer et al. 2015 |
| COL | South America | *M. tuberculata* | AY283078 | Facon et al. 2003 |
| CSM2 | Africa | *M. tuberculata* | KP774658 | Van Bocxlaer et al. 2015 |
| CTA | Africa | *M. tuberculata* | KP774657 | Van Bocxlaer et al. 2015 |
| FAL | Caribbean | *M. tuberculata* | AY283075 | Facon et al. 2003 |
| KIS | Africa | *M. tuberculata* | AY283076 | Facon et al. 2003 |
| KLV | Africa | *M. tuberculata* | AY791931 | Sørensen et al. 2005 |
| Not Determined/LMN | Africa | *M. tuberculata* | AY456616 | Wilson et al. 2004 |
| Not Determined/LM* | Africa | *M. tuberculata* | AY456617 | Wilson et al. 2004 |
| LMI | Africa | *M. tuberculata* | KP774659 | Van Bocxlaer et al. 2015 |
| LMI | Africa | *M. tuberculata* | KP774661 | Van Bocxlaer et al. 2015 |
| LMI | Africa | *M. tuberculata* | KP774662 | Van Bocxlaer et al. 2015 |
| LMI | Africa | *M. tuberculata* | KP774664 | Van Bocxlaer et al. 2015 |
| LMI | Africa | *M. tuberculata* | KP774665 | Van Bocxlaer et al. 2015 |
| LMI | Africa | *M. tuberculata* | KP774666 | Van Bocxlaer et al. 2015 |
| LMI | Africa | *M. tuberculata* | KP774667 | Van Bocxlaer et al. 2015 |
| LMI | Africa | *M. tuberculata* | KP774668 | Van Bocxlaer et al. 2015 |
| LMN | Africa | *M. tuberculata* | KP774660 | Van Bocxlaer et al. 2015 |
| LMN | Africa | *M. tuberculata* | KP774663 | Van Bocxlaer et al. 2015 |
| MAC 1 | Africa | *M. tuberculata* | AY791911 | Sørensen et al. 2005 |
| MAC 1 | Africa | *M. tuberculata* | AY791915 | Sørensen et al. 2005 |
| MAL_H2 | Southeast Asia | *M. tuberculata* | KP284119 | Chiu et al. 2018 |
| MAL_H3 | Southeast Asia | *M. tuberculata* | KP284120 | Chiu et al. 2018 |
| MAL_H4 | Southeast Asia | *M. tuberculata* | KP284121 | Chiu et al. 2018 |
| MAL_H5 | Southeast Asia | *M. tuberculata* | KP284122 | Chiu et al. 2018 |
| MAL_H6 | Southeast Asia | *M. tuberculata* | KP284123 | Chiu et al. 2018 |
| MAL_H7 | Southeast Asia | *M. tuberculata* | KP284124 | Chiu et al. 2018 |
| MAL_H8 | Southeast Asia | *M. tuberculata* | KP284125 | Chiu et al. 2018 |
| MAL_H10 | Southeast Asia | *M. tuberculata* | KP284126 | Chiu et al. 2018 |
| MAL_H11 | Southeast Asia | *M. tuberculata* | KP284127 | Chiu et al. 2018 |
| MAL_H12 | Southeast Asia | *M. tuberculata* | KP284128 | Chiu et al. 2018 |
| MAL_H13 | Southeast Asia | *M. tuberculata* | KP284129 | Chiu et al. 2018 |
| MAL | Southeast Asia | *M. tuberculata* | KP284118 | Chiu et al. 2018 |
| MOF | Africa | *M. tuberculata* | AY283074 | Facon et al. 2003 |
| MOO | Polynesia | *M. tuberculata* | AY283082 | Facon et al. 2003 |
| MOO | Polynesia | *M. tuberculata* | AY283083 | Facon et al. 2003 |
| ND | Asia | *M. tuberculata* | AY283081 | Facon et al. 2003 |
| OMW | Middle-East | *M. tuberculata* | AY283073 | Facon et al. 2003 |
| PDC | Caribbean | *M. tuberculata* | AY283071 | Facon et al. 2003 |
| PHI | Southeast Asia | *M. tuberculata* | AY456618 | Wilson et al. 2004 |
| RLK | Africa | *M. tuberculata* | KP774669 | Van Bocxlaer et al. 2015 |
| Seychelles | Africa | *M. tuberculata* | AY283077 | Facon et al. 2003 |
| TAI | Africa | *M. tuberculata* | AY791910 | Sørensen et al. 2005 |
| TUM | South America | *M. tuberculata* | AY283080 | Facon et al. 2003 |
| ULA | Africa | *M. tuberculata* | AY791930 | Sørensen et al. 2005 |
| ULE | Africa | *M. tuberculata* | KP774672 | Van Bocxlaer et al. 2015 |
| ULK | Africa | *M. tuberculata* | KP774671 | Van Bocxlaer et al. 2015 |
| UNI | Africa | *M. tuberculata* | KP774670 | Van Bocxlaer et al. 2015 |
|  | Caribbean | *M. amabilis* | AY283068 | Facon et al. 2003 |
|  | Africa | *M. cf. liebrechtsi* | KP774641 | Van Bocxlaer et al. 2015 |
|  | Africa | *M. cf. liebrechtsi* | KP774646 | Van Bocxlaer et al. 2015 |
|  | Africa | *M. cf. nsendweensis* | KP774642 | Van Bocxlaer et al. 2015 |
| MAG | Africa | *M. magnifica* | AY791928 | Sørensen et al. 2005 |
| NYA | Africa | *M. nyassana* | AY791933 | Sørensen et al. 2005 |
| NYA/POL | Africa | *M. nyassana/polymorpha* | AY791932 | Sørensen et al. 2005 |
| POL | Africa | *M. polymorpha* | AY791921 | Sørensen et al. 2005 |
| POL | Africa | *M. polymorpha* | AY791923 | Sørensen et al. 2005 |
| POL | Africa | *M. polymorpha* | AY791924 | Sørensen et al. 2005 |
| PUB | Africa | *M. pubiformis* | AY791920 | Sørensen et al. 2005 |
| TRU | Africa | *M. truncatelliformis* | AY791917 | Sørensen et al. 2005 |
| TUR | Africa | *M. turritispira* | AY791908 | Sørensen et al. 2005 |
| MAC 2 | Africa | *M. virgulata* | AY791907 | Sørensen et al. 2005 |
| TAN | Africa | *M. virgulata* | AY791906 | Sørensen et al. 2005 |
| WOO | Africa | *M. woodwardi* | AY791901 | Sørensen et al. 2005 |
| WOO | Africa | *M. woodwardi* | AY791902 | Sørensen et al. 2005 |
| WOO | Africa | *M. woodwardi* | AY791903 | Sørensen et al. 2005 |
| WOO | Africa | *M. woodwardi* | AY791904 | Sørensen et al. 2005 |
| WOO | Africa | *M. woodwardi* | AY791905 | Sørensen et al. 2005 |
| WOO/TRU | Africa | *M. woodwardi/trucatelliformis* | AY791900 | Sørensen et al. 2005 |
|  | Southeast Asia | *Stenomelania sp.* | KU318368 | Ng et al. 2016 |
|  | Not Determined | *T. granifera* | AY283069 | Facon et al. 2003 |
|  | Africa | *Cleopatra bulimoides* | AY791935 | Sørensen et al. 2005 |
|  | Africa | *Lavigeria grandis* | AY456594 | Wilson et al. 2004 |

### Qualitative Morphological Characterization

Previous studies using distinct features of shell morphology, such as color patterns (ornamentation) and sculpture, have characterized phenotypes of *M. tuberculata,* which provides a quick, practical foundation to qualitatively assess differences in phenotypic variation. Thus, we evaluated phenotypic variation in snail shells using distance-based rank-scoring methodologies first employed by Samadi et al. (1999) and later modified in other studies (Facon et al. 2003; Sørensen et al. 2005; Van Bocxlaer et al. 2015). Unfortunately, this progressive modification resulted in inconsistency of measured variables, and consequently, character ranking across studies. Therefore, we did not include those analyses and results in our study as they were not reliable.

Therefore, when building our character matrix, we eliminated previously used qualitative variables that were non-applicable, or ambiguous, from our data. These included various shell colors, conicity of shell, roundness of whirl, and subsutural spiral cord (Facon et al. 2003; Sørensen et al. 2005; Van Bocxlaer et al. 2015). Further, calcium carbonate deposition distorted some of the shell sculptures patterns in most of our individuals. Therefore, we removed the spiral groove, reticulate indentation, and reticulate ridge characters from our analyses. We retained the axial rib character, because it was pronounced despite distortions caused by deposition. We removed snails from morphological analysis if deposition prevented accurate assessment of any character. Accordingly, we scored a suite of 11 morphological characters (Table S2). However, we still present all shell sculpture patterns to document our observations (Figure 4A).

We determined ranks for qualitative variables using dissimilarity-based systematics and implemented variable rankings in a manner that sought to maximize dissimilarity by maximizing the rank distance between dissimilar characters. For example, background shell coloration varied from very pale to dark, thus a rank of 1 was assigned to pale and 4 for dark. Rank scores for all qualitative variables were Z-transformed to stabilize the variance among disproportionate variable scores. We then used the transformed rank scores to construct a distance matrix that we ordinated using Non-Metric Multi-Dimensional Scaling (nMDS) analyses using 11 dimensions and 1000 iterations. Because not all snails exhibited all of the qualitative characters, we used a Euclidean distance method that accounts for missing values. We performed these analyses using R version 3.3.1 and packages *vegan v.2.4-1* and *MASS 7.3-45.*

## Results

### Haplotype Composition and Distribution

Three unique 16S rRNA gene haplotypes were present in snails (*n* = 165) collected in the CR, SMR, and GR (GenBank Accession: XXXXXXX-XXXXXXX). We arbitrarily labeled each haplotype H1, H2, and H3. In a median-joining network, H1 is 32 mutational steps from the H2 haplotype and 34 steps from H3 (Figure 2A). The H2 and H3 haplotypes differ by five mutational steps. H1 was the most abundant haplotype (*n* = 108) and occurred across multiple sampling localities. However, in the SMR, H1 was not detected below the SMR5 site (Figure 2C). In the GR, H1 was found at a site directly below the outflow of a diverted channel (Figure 2E). The second most abundant haplotype was H2 (*n* = 54), which was found in the SMR between sites SMR8 and SMR14 (Figure 2C). One individual carrying the H2 haplotype was found in the CR River (site Old Channel; Figure 2D) and six individuals were found in the GR (site AbCH1; Figure 2E). H3 was the rarest haplotype (*n* = 3) and was only detected at the Old Channel site from the CR (Figure 2D).

**Figure 2.**
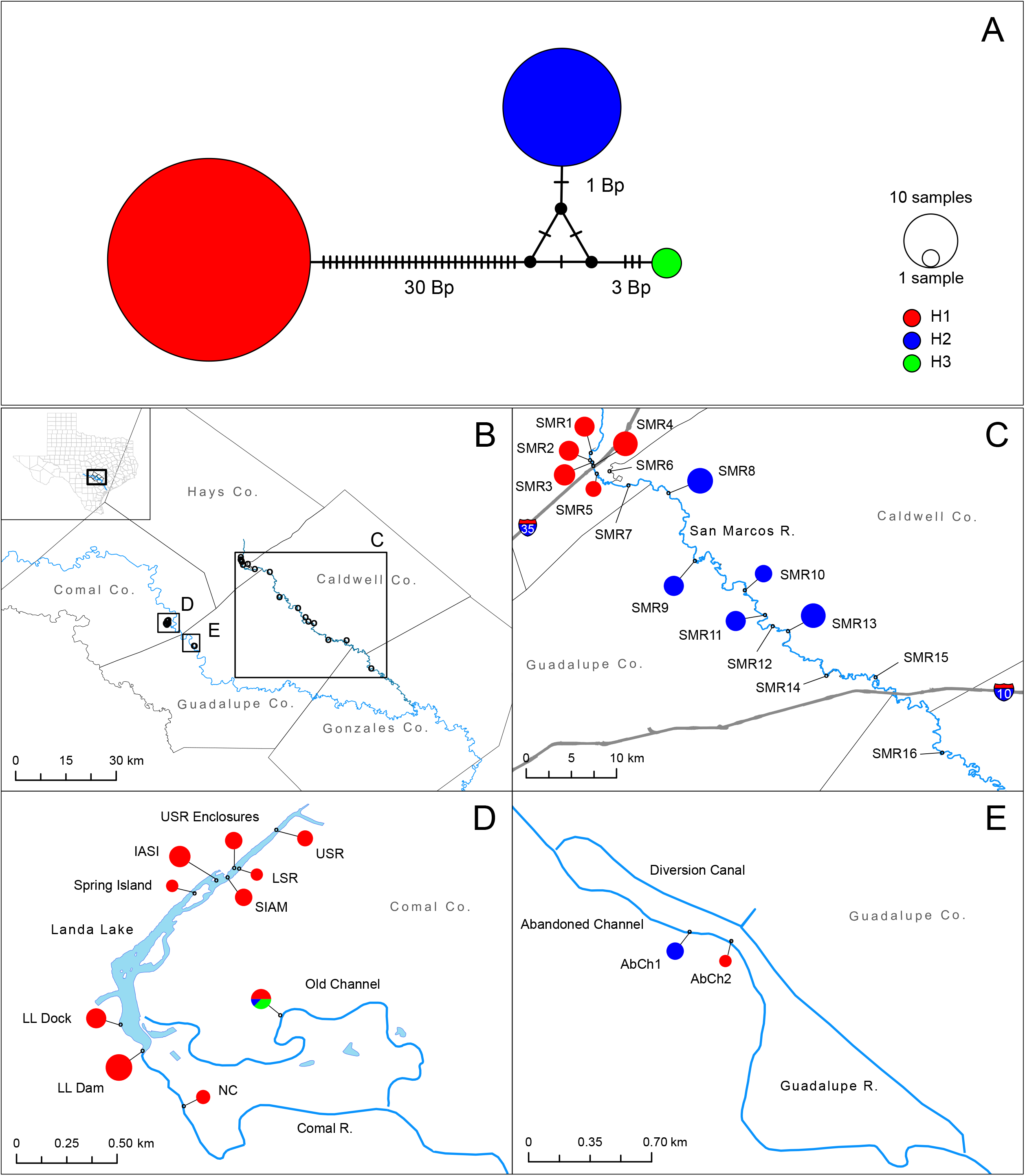
(A) Median-joining haplotype network of 16S rRNA sequences from *Melanoides tuberculata* collected in central Texas (Red = H1, Blue = H2, and Green = H3). The number of mutational steps in base pairs (bp) for all three haplotypes are denoted by hashes with the number of changes from next closest haplotype. Closed black circles represent hypothetical haplotypes between each haplogroup. The size of the colored circles is scaled to the relative frequency of each haplotype. (B) Map of the sampled rivers where *M. tuberculata* were collected (sample site = open black circles). (C) San Marcos River; (D) Comal River; and (E) Guadalupe River. Each solid circle is a pie chart showing the composition of haplotypes collected at each site sampled. The size of the chart shows the relative sample size of snails collected.

### Phylogenetic Analyses

MODELTEST returned HKY85+Γ as the best fit model for the 16S rRNA gene sequence data. Bayesian phylogenetic analyses of *M. tuberculata* from central Texas and reference sequences available on GenBank data returned nine major clades (Figure 3). The outgroup *Stenomelania* sp. was placed within its own clade, and the same was true for the other outgroup taxa; *M.* cf. *nsendweensis, M.* cf. *liebrechtsi,* and *T. granifera.* The *M. polymorpha* complex proposed by Sørensen et al. (2005) formed another clade. We only assigned numbers to those clades containing *M. tuberculata*. Clade 1 contains one *M. tuberculata* morph with an undetermined region of origin (AY283081) and *M. amabilis*. Clade 2 contains several *M. tuberculata* morphs (BCI, CAE, all CDI, all LMI, MAC 1, TUM, MAL, MAL H2-H5, and MAL H11), the Texas H1 haplogroup, and *M. virgulata* morphs (TAN and MAC 2). Clade 3 contains the Polynesian morphs MOO 1 and MOO 2. Clade 4 contains four possible sub-clades; however, sub-clade 4.5 has low bootstrap support. All morphs within this clade are identified as *M. tuberculata.* Sub-clade 4.1 contains morphs BIT, COL, PHI, FAL, MAL H6, MAL H8, MAL 12, MAL H13, and the Texas H3 haplogroup. Sub-clade 4.2 is represented by a single morph (MAL H10). Sub-clade 4.3 contains morphs PDC and MAL H7. Sub-clade 4.4 contains Seychelles, TAI, LMN, KIS and BOU. Sub-clade 4.5 contains MOF, CHO, CLE, CSM2, CTA, KLV, OMW, RLK, ULA, ULE, ULK, UNI, and the Texas H2 haplogroup. Our phylogenetic reconstruction supports previously published results that show *M. tuberculata* is polyphyletic and shows high diversity at the 16S rRNA gene (Facon et al. 2003; Sørensen et al. 2005; Van Bocxlaer et al. 2015).

**Figure 3.**
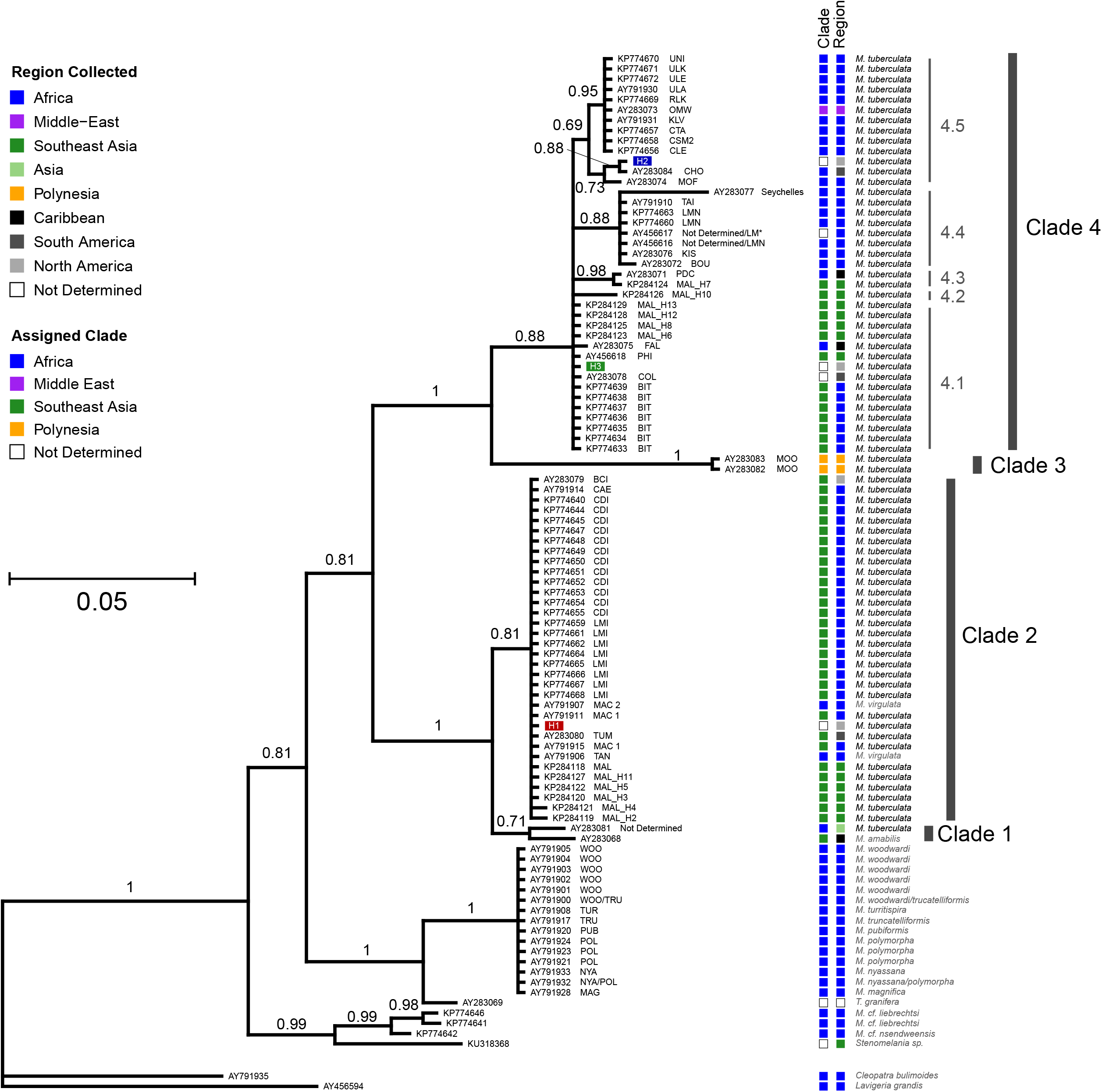
Phylogeny of *Melanoides* spp. based on Bayesian analysis of the 16S rRNA mitochondrial gene (250 bp). Posterior probabilities are given for internal nodes, and GenBank accession numbers, morph codes, and species names are shown for terminal taxa. The scale bar represents the number of base pair changes per site. In the legend, Region Collected represents the region where each snail was collected, and Assigned Clade depicts geographic origins inferred from previous phylogenetic analyses. Haplotypes collected in Texas are highlighted by colored boxes. Major and minor clades are labeled and identified with grey bars.

### Morphological Characterization

We identified 62 unique phenotypes based on 11 qualitative variables (Table S2). The differences in qualitative rank scores among the haplotypes were visualized in a nMDS ordination plot (Figure 4A). When morphological characters were associated with haplogroup, H2 displays phenotypic separation from the other two haplotypes. We observed seven phenotypes within haplogroup H2 (Table S2). H2 can be qualitatively characterized as having translucent tan shells, lacking dense and deep axial ribs, and displaying thinner ornamentation patterns. They typically do not have a columellar band, but it is very thin when present. We observed three phenotypes within haplogroup H3 (Table S2). H3 is nested inside a cluster that contains several H1 phenotypic variants. H3 can be characterized as having a translucent brown – red shell, lacking dense and deep axial ribs, having thick columellar bands, and exhibiting larger ornamentation patterns compared to haplogroup H2. We observed 51 phenotypes within haplogroup H1 (Table S2); thus, it has the most phenotypic variability among the three haplogroup sampled (Figure 4B). Qualitatively, haplogroup H1 can be characterized as having wide variability in ornamentation patterns, axial rib morphology, and shell background color and intensity; however, adults do not exhibit translucent shells. The accessory datasets generated for this study are available in the Dryad Digital Repository (DRYAD LINK).

**Figure 4.**
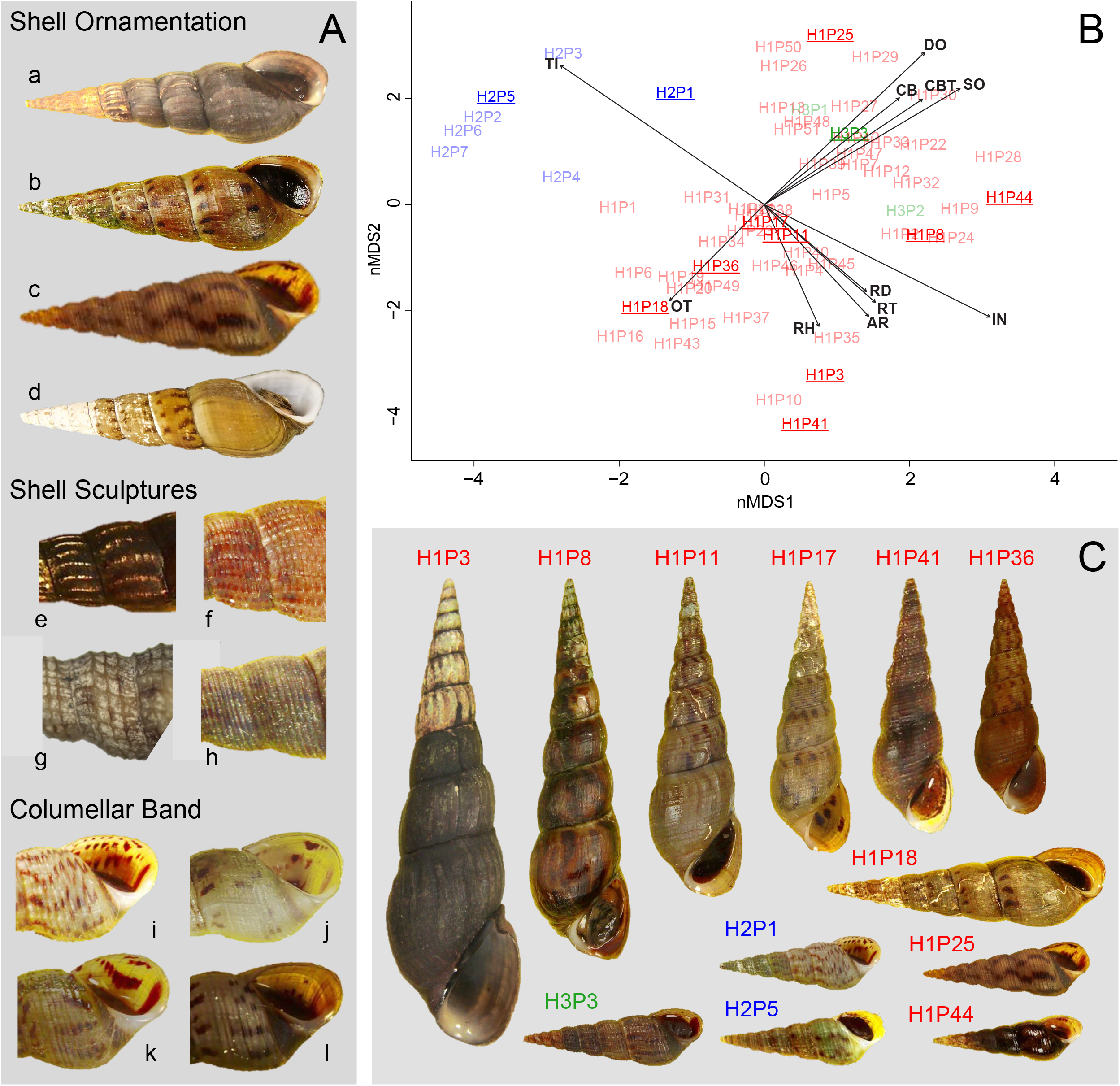
(A) Photographic examples of qualitative characters used to describe shell morphology of *Melanoides tuberculata* found in central Texas. Shell ornamentation is as follows: a) none, b) spots along the suture, c) flames, and d) flames over spots. Shell Sculptures examples are: e) axial ribs, f) reticulate ridge, g) reticulate indentation, and h) spiral grooves. Columellar Band examples are: i) narrow, j) diffuse, k) medium, and l) wide. (B) Plotted output of nMDS analysis of qualitative characters. Data points are point estimates of phenotypes and associated with 16S haplotype. Each phenotype (n = 62) is colored by their respective haplotype, and those in bold are depicted in panel C. Vectors are labeled with character code and represent the strength of dissimilarity among the qualitative variables. (C) Subset of *M. tuberculata* phenotypes used in nMDS analysis. Snails are labeled by haplogroup and designated phenotype (see Table S2). Shell sculptures f, g, and h were shown to document the observation of these characters in some shells in good condition; although, they were not included our phenotypic assessments (see Methods and Discussion).

## Discussion

We have described genetic, phenotypic, and ecological variation among *Melanoides tuberculata sensu lato* from three central Texas rivers, the suspected point of introduction into North America based on temporal sampling (Murray 1964; Murray and Wopschall 1965; Murray 1971). At our sampling sites, these snails are primarily parthenogenetic and disproportionately female (male abundance = 1.14%; Table S1); thus, we expected to detect a single clonal lineage if one introduction event occurred in the past. However, our results show at least three unique haplotypes of *Melanoides* sp. occur in central Texas based on 16S rRNA sequence data.

Our results demonstrated an association between haplotype geographic distribution and water temperature profiles of sites. Specifically, H1 was the dominant haplotype in the CR and the only haplotype detected in headwaters of the SMR (SMR1-SMR5). This is significant because the headwaters of the SMR and the CR are spring systems with stable water temperatures (Crowe and Sharp Jr 1997; Saunders et al. 2001). We showed temperature fluctuation for these areas did not exceed the experimentally determined lethal limit proposed by Mitchell et al. (2005) during the course of our study, and they are consistent with published aquatic temperature profiles (Crowe and Sharp Jr 1997; Saunders et al. 2001). Therefore, our study suggests H1 is limited to waters with stable temperatures that do not exceed 18°C in central Texas.

In the SMR, H2 was the only haplotype found below site SMR5. This is significant because a previous evaluation of spring flows in the SMR demonstrated the river has a different temperature profile after the SMR7 site (Saunders et al. 2001). Specifically, water temperatures below SMR 7 are more variable and water temperatures seasonally drop below 18°C. The geographic distribution of each haplogroup shows clear partitioning in the SMR suggesting there are differences in habitat preferences among these genetic variants (Figure 2C). In the CR, H2 was only detected at the Old Channel site, and only one snail carrying the H2 haplotype was collected during our surveys (Table 2), which indicates that H2 snails are cold water tolerant while H1 snails are not.

Additional sampling is still needed along the GR; however, partitioning is clearly observed at the two sites we sampled. Haplotype H1 was only found at the AbCH2 site and H2 was only found upstream at the AbCH1 site. Our water temperature data from AbCH1 showed water temperature fluctuated between 10.5° and 32.4°C (Table 1). Even though limited in duration, water temperature data for the AbCH2 shows temperatures did not fall below 18°C at this site, while the upstream AbCH1 logger showed temperatures fell below 18°C during the same time frame (Figure S1). It is important to note the AbCH2 site is located directly below the outflow of a diverted channel; therefore, we speculate this site has some groundwater input and serves as a seasonal refuge for snails, which is supported by our water temperature data.

By using mitochondrial DNA sequence data, we have determined the likely origin of each haplogroup of *M. tuberculata* found in central Texas. We constructed a phylogeny using all available sequence data, including the three haplogroups detected in Texas (H1, H2, and H3), and we inferred global morphs could be partitioned into five major clades (Figure 3). Haplotype H1 nested within Clade 2. Many of the snails within Clade 2 have been identified as invasive lineages and are likely of Asian origin (Facon et al. 2003; Genner et al. 2004; Van Bocxlaer et al. 2015). The phylogenetic placement of Malaysian haplogroups (Chiu et al. 2018) with these invasive lineages supports these conclusions; thus, the Texas haplogroup H1, as well as all of the morphs within Clade 2, likely originated in Southeast Asia.

H2 falls within Clade 4, which is a mixture of morphs found in the Middle East and Africa, and it also includes invasive lineages from Southeast Asia. Except for the OMW morph, which was collected in the Middle East and is most likely an invasive there, most morphs within sub-clade 4.5 are likely from Africa (Facon et al. 2003; Genner et al. 2004; Van Bocxlaer et al. 2015). Haplotype H2 detected in Texas is identical to the sequence of invasive morph CHO (Venezuela), which is also clustered within sub-clade 4.5. Thus, we speculate that haplogroup H2 is originally from Africa but may have been transported to South America via North America or vice versa. Inferring the specific origin of haplogroup H3 is more complex, because it is included in Sub-clade 4.1, which is comprised of several invasive morphs (COL, FAL, PHI, and BIT 1-7) that are most likely from Southeast Asia (Facon et al. 2003; Van Bocxlaer et al. 2015). This is supported by the placement of several Malaysian morphs within the same sub-clade. Facon et al. (2003) were unable to determine an origin of the COL morph, because it was not genetically similar to any other morph in that study, which included snails from the Caribbean, the Americas, Europe, Africa, the Middle East, Asia and Polynesia. However, subsequent phylogenetic analyses of 16S sequences placed COL in a clade with FAL and an invasive morph labeled ND (not determined) from the Philippines (Sørensen et al. 2005). Our NJ distance matrix showed COL has 99.59% sequence identity with FAL and 100% identity with the ND sample (AY456618), which we labeled as PHI to resolve the ND designation. Furthermore, we were able to incorporate several Malaysian haplotypes (MAL H6-H8, MAL H12, and MAL H13; Chiu et al. 2018) deposited into GenBank in our analyses, which may not have been available to previous studies (Table 3). Therefore, it is likely that all the morphs within Sub-clade 4.1 are from Southeast Asia, and specifically, haplogroup H3 detected Texas is originally from the Philippines.

Our haplotype network demonstrates the genetic distance in base pair differences (i.e. mutations) among each of the haplotypes detected in Texas. The 16S rRNA gene contains regions that are highly conserved across taxonomic groups, but it also contains regions with higher mutation rates (Coenye and Vandamme 2003; Yang et al. 2014). For these reasons, it is used to identify eukaryotes to the species level. Estimated mutation rates at the 16S rRNA locus for many eukaryotes remain elusive; however, among those examined, current estimates of the average base pair mutation rate across the mitochondrial genome ranges from 1.37 × 10^-7^ to 12.2 × 10^-9^ per site per generation (Denver et al. 2000; Haag-Liautard et al. 2008; Lynch et al. 2008; Xu et al. 2011). Six months is the estimated generation time for *M. tuberculata* (Livshits and Fishelson 1983). If we applied the fastest eukaryotic mitochondrial genome mutation rate, then we estimated it would take an average of ~3.125 myr to accumulate one bp mutation in a population of snails. Given the timeline of invasion and the magnitude of mutational steps between the haplotypes, it is reasonable to assume these haplotypes represent separate introduction events and not evolution within our system. Although, our estimates should be viewed with caution owing to uncertainty of the mutation rate within *Melanoides*.

Furthermore, we can confidently assume H3 and H2 represent separate introductions and not evolution, because H3 was not detected in the SMR. If we assume evolution, given the relative frequency of H2 in the lower SMR versus the CR, then we would expect to find H3 in the SMR where H2 is most abundant, however, we did not. With respect to similar haplotype composition across the rivers (i.e., the presence of H1 and H2 in all three rivers surveyed), we speculate this can be explained by their close proximity to each other. Additionally, all three rivers are popular recreational areas, and snails could have been easily transported between them through shared user traffic (Mitchell et al. 2007).

The breadth of phenotypic variation observed in haplogroup H2 is much narrower than the variation observed in H1. Specifically, H2 contained seven phenotypes and H1 contained 51 phenotypes (Table S2). The primary distinguishing characteristics of the H2 haplogroup in central Texas are a translucent tan shell, a general lack of a columellar band, and general lack of axial ribs. If the columellar band is present, it is very thin and diffuse. Axial ribs are mild in thickness and shallow in depth if they are present. Additionally, axial ribs are distributed heterogeneously among whorls, tend to be located towards the apex of the shell, and do not progress to the body whorl. Even though we did not use additional shell sculpturing as diagnostic characters, H2 individuals did exhibit reticulate indentations on the whorls (Figure 4A).

We cannot draw any conclusions regarding the breadth of phenotypic variation with the H3 haplotype owing to small sample size. Regardless, among those H3 individuals collected, they exhibited a translucent shell with either an amber or a darker copper background shell color, a dense flame ornamentation pattern, a thick columellar band, and reticulate ridge sculpturing patterns (Figure 4A). Additional sampling is needed to better estimate the phenotypic variation in H3; although, they are relatively rare in our sampling area.

Comparatively, the H1 haplogroup has broad phenotypic variation with many defined morph categories; however, the primary distinguishing characteristics that separates it from haplogroups H2 and H3 is the lack of translucent shells and darker shell color in the adults. Juveniles will have translucent shells but usually with darker shell coloration when compared to H2 snails, pronounced columellar bands, and thick/robust shell ornamentation patterns. We also acknowledge that omission of shell sculpture patterns may have reduced the discriminatory power of the nMDS analysis. We speculate H3 could be distinguished from H1 using the reticulate ridge and spiral groove characters; however, we took a conservative approach owing to distortion of the shell by mineral deposition.

Qualitative analyses of morphological variation support phenotypic differentiation of the H2 haplogroup from the H3 and H1 haplogroups. However, haplogroups H1 and H3 have phenotypes with similar shell color and ornamentation patterns (Figure 4B). H3 can be differentiated from H1 only by subtle differences in shell morphology such as reticulate ridge sculptures and shell color. Therefore, phenotypic similarities between H1 and H3 may confound their identification in the field.

Another approach that could possibly discriminate among morphs is quantitative morphology (i.e., morphometrics). However, morphometric approaches for differentiating wild snails are complicated by shell plasticity under varying environmental conditions (Samadi et al. 2000; Gustafson et al. 2014; Abdelhady et al. 2018; Albarrán-Mélzer et al. 2019). Further, common garden experimentation has demonstrated that characters based on shell geometry are unreliable as a diagnosis of gastropod taxa collected from the wild (Gustafson et al. 2014; Albarrán-Mélzer et al. 2019). Therefore, these types of analyses should be conducted when environmental variation can be controlled.

Consequently, environmentally induced phenotypic plasticity as well as inherent variability in experience and subjectivity among assessors may lead to misidentifications. Therefore, our study supports previous suggestions that a combined genetic, morphological, and ecological approach should be used to identify morphs of *Melanoides tuberculata sensu lato* rather than relying solely on morphology (Facon et al. 2003; Genner et al. 2004; Van Bocxlaer et al. 2015). Accordingly, we recommend common garden experimentation on Texas haplotypes to produce a more rigorous phenotypic characterization as it relates to plasticity.

*Melanoides tuberculata* appear to possess robust physiological tolerances that are not yet fully characterized. Some known examples include tolerance to saline waters (Russo 1973; Roessler et al. 1977), resistance to desiccation (Facon et al. 2004), and tolerance to decontaminants (Mitchell et al. 2007). Our phylogenetic analyses support different geographic origins among Texas *Melanoides*, and we have shown physiological tolerances and environmental preferences vary across different haplotypes in natural systems. Taken together, these observations are evidence that *M. tuberculata sensu lato* varies in the ability to colonize different habitats and concomitantly their potential for spread. Therefore, the observed habitat partitioning among the haplotypes within the SMR may be due to advantageous physiological or behavioral adaptations of H2 compared to the H1 haplotype, but that is yet to be explored.

However, the role of inter-haplotype competition is unclear. We cannot dismiss the possibility that H1 may be competitively excluding H2 from warmer waters. This may be acting synergistically with the cold-water tolerance observed in H2, and thus is another explanatory factor contributing to haplotype partitioning in the SMR. Evidence for this is given in the haplotype composition at the Old Channel site in the CR. This site exhibits stable water temperatures and the only one to contain multiple haplotypes; however, H1 was still the most abundant (Figure 2D).

Additional monitoring and genetic testing should continue throughout the region, because *Melanoides* snails are first intermediate hosts for multiple species of digenetic trematode parasites. Several snail parasites are the cause of major human health problems in Asia (Pinto and De Melo 2011). In particular, four exotic trematode species: *Haplorchis pumilio, Centrocestus formosanus*, *Philophthalmus gralli*, and a recently detected cercaria presumed to be a *Renicola* sp. presently occur in Texas, which is a direct consequence of the establishment of their snail hosts (Nollen and Murray 1978; D.G. Huffman, personal communication 2016; Fleming et al. 2011; Pinto and De Melo 2011). These trematodes have a high level of host-specificity for their first intermediate snail host, and therefore, cannot complete their life cycles beyond the range of *M. tuberculata*. Parasites of *Melanoides tuberculata sensu lato* might follow their snail hosts into new habitats that were thought to be protected by previously reported thermal limits of the snails. Our data show that this is not the case when thermal tolerance varies.

While managers are most interested in the deleterious effects introduced parasites may have on native vertebrates, such as the federally listed fountain darter *(Etheostoma fonticola)* and devils river minnow (*Dionda diaboli*), an alternative approach to parasite mitigation might be managing their thiarid snail hosts. Now that we have disentangled morphological, genetic, and ecological variation among *M. tuberculata* in central Texas, controlled experiments investigating differences among these haplotypes are now possible. These experiments should test differences in adaptive behaviors, thermal tolerances, differential suitability as hosts for parasites, and other life history traits.

Our results have shown that central Texas populations of *M. tuberculata sensu lato* have a wider distribution than previously described, in both geography and habitat preferences. This breadth is owing to cryptic diversity and multiple introduction events of at least three different lineages. Each of these likely have different adaptive strategies that evolved in their native ranges, highlighting the need to address and manage three invasive taxa instead of one. Our results also agree with other genetic studies showing *M. tuberculata* is polyphyletic and in need of taxonomic revision (Facon et al. 2003; Van Bocxlaer et al. 2015). Further, despite mostly using parthenogenesis as reproductive strategy, *Melanoides* sp. can also reproduce sexually and bear viable offspring when males are present (Samadi et al. 1999). Studies of hybrids resulting from cross-clonal breeding suggest hybrids outcompete and hold selective advantages over their parental morphotypes (Facon et al. 2005); thus, multilocus genotyping in this and other invaded environments can confirm whether hybridization is another characteristic that will facilitate additional invasions and spread. In this study, we have found invasion dynamics and invasive potential of different lineages of *Melanoides* sp. are concealed by remarkable phenotypic plasticity, which can only be parsed via genetic identification.

## Acknowledgements

We would like to thank Evan Valenta, Mclean Worsham, James D. Evans, Cassandra P. Mccumber-Aguilar, Angel M. Peltola, Carlos Baca, Jeremiah Leach, Dakota L. Rhoad, Amanda C. Andrews, Gabriella D. Solis, Thomas Marshall, Mireya Escandon, Nick Porter and Jube Guajardo for assistance with data collection. This research was supported by BIO-West, Inc., the San Marcos Aquatic Resources Center, and start-up funds from Texas State University to DR. We also thank Mr. CW Grumbles for property access at site SMR9.

## SUPPLEMENTARY MATRIALS

**Table S1.**
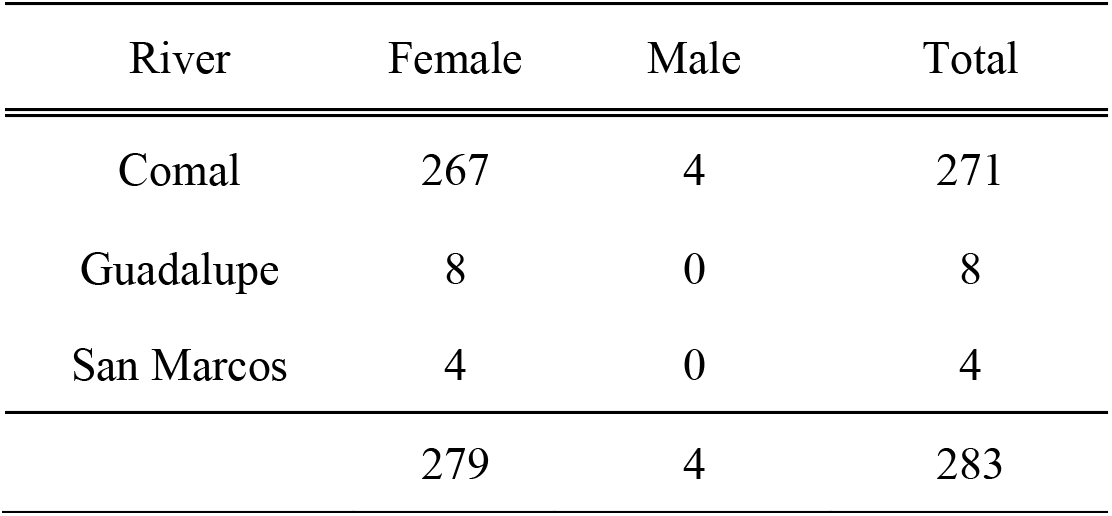
Summary of female and male abundance of *Melanoides tuberculata* surveyed in the Comal, Guadalupe, and San Marcos rivers. Sex was determined by visually identifying sperm in the digestive gland of the snails.

**Table S2.** *Melanoides tuberculata* modified qualitative data matrix showing the ranks of 11 shell characteristics for each unique phenotype (P# = arbitrary phenotype identifier), the river system it was collected from (CR = Comal River, TX; SMR = San Marcos River, TX), and the 16S haplotype designation.

| Phenotype | River | Haplotype | Background Color |  | Ornamentation |  |  | Columellar Band |  | Sculptures |  |  |  |
| --- | --- | --- | --- | --- | --- | --- | --- | --- | --- | --- | --- | --- | --- |
|  |  |  | IN | TI | DO | OT | SO | CB | CBT | AR | RH | RT | RD |
| P1 | CR | H1 | 2 | 3 | 2 | 3 | 1 | 0 | * | 0 | * | * | * |
| P2 | CR | H1 | 4 | 1 | 3 | 2 | 2 | 0 | * | 1 | 1 | 2 | 3 |
| P3 | CR | H1 | 4 | 1 | 0 | * | * | 0 | * | 1 | 1 | 2 | 3 |
| P4 | CR | H1 | 4 | 3 | 2 | 2 | 2 | 0 | * | 1 | 2 | 1 | 1 |
| P5 | CR | H1 | 3 | 3 | 3 | 3 | 2 | 1 | 3 | 1 | 2 | 1 | 1 |
| P6 | CR | H1 | 3 | 3 | 1 | 3 | 1 | 0 | * | 0 | * | * | * |
| P7 | CR | H1 | 3 | 3 | 3 | 1 | 2 | 1 | 4 | 1 | 2 | 1 | 1 |
| P8 | CR | H1 | 4 | 2 | 3 | 1 | 3 | 0 | * | 1 | 2 | 1 | 2 |
| P9 | CR | H1 | 4 | 2 | 3 | 2 | 3 | 1 | 4 | 1 | 2 | 1 | 1 |
| P10 | CR | H1 | 4 | 1 | 1 | 5 | 1 | 0 | * | 1 | 1 | 1 | 2 |
| P11 | CR | H1 | 3 | 3 | 2 | 2 | 2 | 0 | * | 1 | 1 | 1 | 2 |
| P12 | CR | H1 | 3 | 3 | 3 | 2 | 3 | 1 | 1 | 1 | 1 | 1 | 2 |
| P13 | CR | H1 | 2 | 3 | 3 | 2 | 3 | 0 | * | 0 | * | * | * |
| P14 | CR | H1 | 3 | 3 | 1 | 2 | 3 | 0 | * | 0 | * | * | * |
| P15 | CR | H1 | 3 | 3 | 0 | * | * | 0 | * | 1 | 2 | 2 | 3 |
| P16 | CR | H1 | 2 | 3 | 1 | 5 | 1 | 0 | * | 1 | 2 | 1 | 1 |
| P17 | CR | H1 | 2 | 3 | 3 | 5 | 2 | 1 | 2 | 1 | 2 | 1 | 1 |
| P18 | CR | H1 | 2 | 3 | 2 | 5 | 1 | 0 | * | 1 | 2 | 1 | 1 |
| P19 | CR | H1 | 2 | 3 | 2 | 5 | 1 | 1 | 1 | 1 | 2 | 1 | 1 |
| P20 | CR | H1 | 2 | 3 | 2 | 5 | 2 | 0 | * | 1 | 2 | 1 | 1 |
| P21 | CR | H1 | 2 | 3 | 2 | 3 | 2 | 1 | 2 | 1 | 2 | 1 | 1 |
| P22 | CR | H1 | 3 | 3 | 3 | 1 | 3 | 1 | 4 | 1 | 1 | 1 | 1 |
| P23 | CR | H1 | 2 | 3 | 3 | 5 | 2 | 1 | 1 | 1 | 2 | 1 | 1 |
| P24 | CR | H1 | 4 | 1 | 3 | 2 | 3 | 0 | * | 1 | 1 | 2 | 2 |
| P25 | SMR | H1 | 2 | 4 | 3 | 1 | 3 | 1 | 4 | 0 | * | * | * |
| P26 | SMR | H1 | 2 | 4 | 3 | 2 | 3 | 1 | 1 | 0 | * | * | * |
| P27 | SMR | H1 | 2 | 4 | 3 | 1 | 3 | 1 | 3 | 1 | 1 | 1 | 1 |
| P28 | SMR | H1 | 4 | 1 | 3 | 2 | 3 | 1 | 4 | 0 | * | * | * |
| P29 | SMR | H1 | 3 | 4 | 3 | 1 | 3 | 1 | 4 | 0 | * | * | * |
| P30 | SMR | H1 | 3 | 2 | 3 | 1 | 3 | 1 | 4 | 0 | * | * | * |
| P31 | SMR | H1 | 2 | 4 | 2 | 3 | 1 | 1 | 2 | 1 | 1 | 1 | 2 |
| P32 | SMR | H1 | 3 | 2 | 3 | 2 | 3 | 1 | 3 | 1 | 2 | 1 | 1 |
| P33 | SMR | H1 | 3 | 4 | 3 | 2 | 3 | 1 | 4 | 1 | 1 | 1 | 1 |
| P34 | SMR | H1 | 3 | 4 | 1 | 2 | 1 | 1 | 4 | 1 | 2 | 1 | 1 |
| P35 | SMR | H1 | 4 | 1 | 2 | 5 | 1 | 1 | 3 | 1 | 2 | 1 | 1 |
| P36 | SMR | H1 | 3 | 3 | 2 | 2 | 1 | 0 | * | 1 | 2 | 1 | 1 |
| P37 | SMR | H1 | 3 | 2 | 1 | 3 | 2 | 0 | * | 1 | 2 | 1 | 1 |
| P38 | SMR | H1 | 3 | 2 | 2 | 2 | 2 | 0 | * | 0 | * | * | * |
| P39 | SMR | H1 | 3 | 4 | 3 | 2 | 2 | 1 | 4 | 1 | 2 | 1 | 1 |
| P40 | SMR | H1 | 3 | 2 | 2 | 2 | 2 | 0 | * | 1 | 1 | 1 | 1 |
| P41 | SMR | H1 | 4 | 1 | 0 | * | * | 0 | * | 1 | 2 | 1 | 1 |
| P42 | SMR | H1 | 2 | 3 | 3 | 1 | 3 | 1 | 1 | 1 | 1 | 1 | 1 |
| P43 | SMR | H1 | 3 | 3 | 1 | 5 | 1 | 0 | * | 1 | 1 | 1 | 1 |
| P44 | SMR | H1 | 4 | 1 | 3 | 1 | 3 | 1 | 3 | 1 | 1 | 1 | 1 |
| P45 | SMR | H1 | 4 | 2 | 2 | 2 | 1 | 1 | 1 | 1 | 1 | 1 | 1 |
| P46 | SMR | H1 | 3 | 3 | 2 | 5 | 2 | 1 | 1 | 1 | 1 | 1 | 1 |
| P47 | SMR | H1 | 3 | 4 | 3 | 2 | 3 | 1 | 1 | 1 | 1 | 1 | 1 |
| P48 | SMR | H1 | 3 | 3 | 3 | 2 | 2 | 1 | 1 | 0 | * | * | * |
| P49 | SMR | H1 | 4 | 3 | 1 | 5 | 1 | 1 | 2 | 0 | * | * | * |
| P50 | SMR | H1 | 2 | 4 | 3 | 1 | 2 | 1 | 4 | 0 | * | * | * |
| P51 | SMR | H1 | 2 | 4 | 3 | 2 | 2 | 1 | 3 | 1 | 1 | 1 | 1 |
| P1 | CR | H2 | 1 | 6 | 3 | 1 | 1 | 1 | 2 | 1 | 1 | 1 | 1 |
| P2 | SMR | H2 | 1 | 6 | 2 | 3 | 1 | 0 | * | 0 | * | * | * |
| P3 | SMR | H2 | 1 | 6 | 2 | 1 | 1 | 1 | 1 | 0 | * | * | * |
| P4 | SMR | H2 | 1 | 6 | 2 | 2 | 1 | 0 | * | 1 | 2 | 1 | 1 |
| P5 | SMR | H2 | 1 | 6 | 2 | 2 | 1 | 0 | * | 0 | * | * | * |
| P6 | SMR | H2 | 1 | 6 | 1 | 2 | 1 | 0 | * | 0 | * | * | * |
| P7 | SMR | H2 | 1 | 6 | 1 | 3 | 1 | 0 | * | 0 | * | * | * |
| P1 | CR | H3 | 3 | 5 | 3 | 2 | 2 | 1 | 4 | 1 | 1 | 2 | 3 |
| P2 | CR | H3 | 4 | 1 | 3 | 2 | 1 | 1 | 4 | 1 | 1 | 2 | 3 |
| P3 | CR | H3 | 4 | 5 | 3 | 2 | 2 | 1 | 4 | 1 | 1 | 2 | 3 |

IN = Intensity of shell background color: (1) Very pale; (2) Pale; (3) Medium; (4) Dark
TI = Background shell color: (1) Dark Brown; (2) Red – Brown; (3) Brown; (4) Amber; (5) Translucent Copper; (6) Translucent Tan
DO = Shell Ornamentation Density: (0) None; (1) Sparse; (2) Medium; (3) Dense
OT = Type of Shell Ornamentation: (1) Flames only; (2) Flame near apex; (3) More Flames than Spots; (4) More Spots than Flames; (5) Spots along the suture
SO = Relative Size of Ornamentation: (1) Small/Narrow; (2) Medium; (3) Large/Wide
CB = Presence of a Columellar Band: (0) No; (1) Yes
CBT = Thickness of Columellar Band (if present): (1) Diffuse; (2) Narrow; (3) Medium; (4) Wide
AR = Presence of Axial Ribs: (0) Absent; (1) Present
RH = Axial Rib Heterogeneity: (1) Homogeneous; (2) Heterogeneous
RT = Thickness of Axial Rib: (1) Very mild; (2) Mild; (3) Prominent
RD = Depth of Axial Rib: (1) Shallow; (2) Medium; (3) Deep
\* = Character was not exhibited

**Figure S1.**
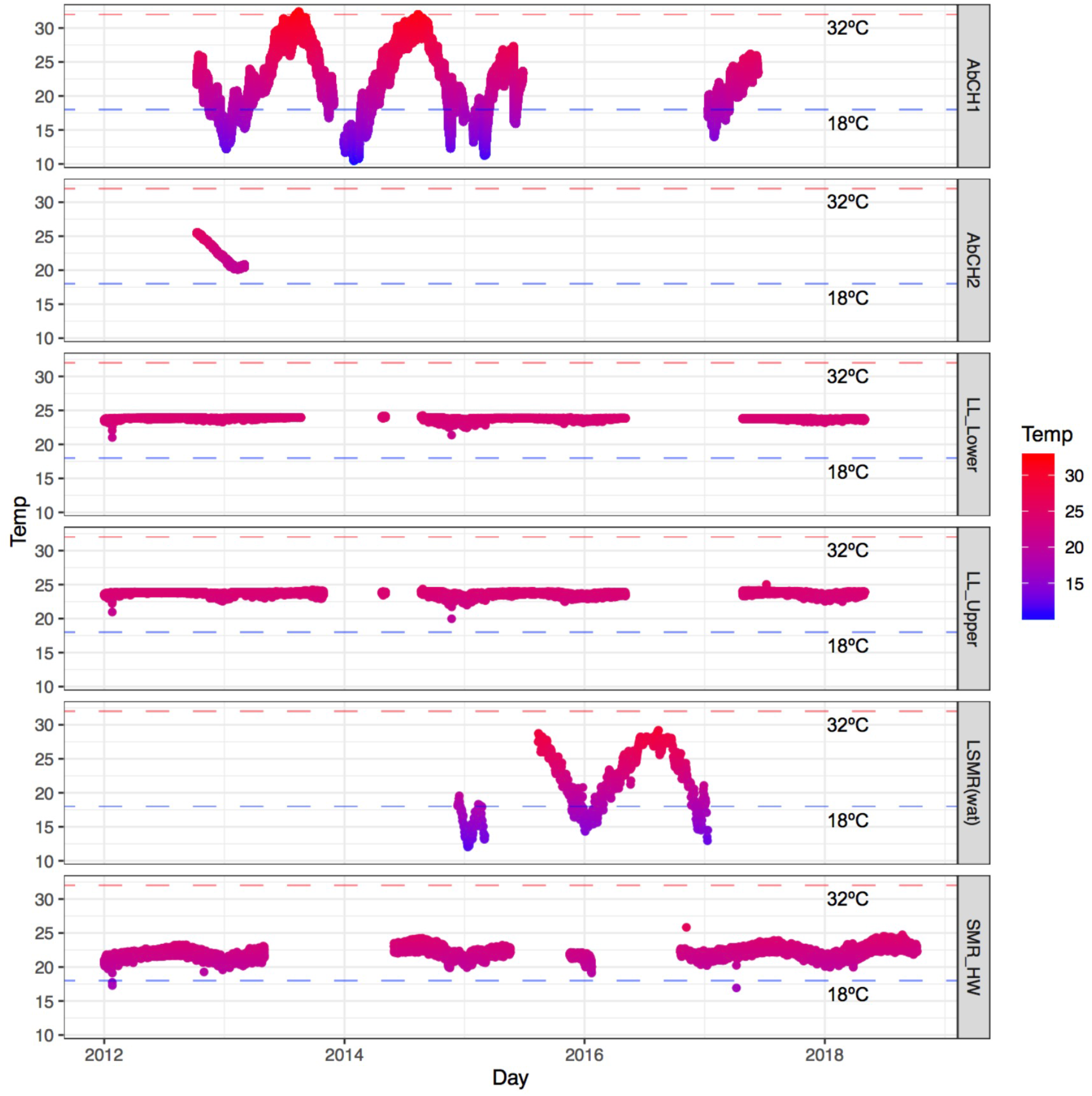
Seasonal ambient temperatures at six sites within the lower San Marcos, Comal, and Guadalupe Rivers in central Texas. Data were collected from January 2012 to October 2018. The critical thermal maximum (32°C) and critical thermal minimum (18°C) for *Melanoides tuberculata* are denoted by red and blue dashed lines, respectively (Mitchell & Brandt 2005). AbCH1 = Abandoned Channel site 1 Guadalupe River, AbCH2 = Abandoned Channel site 2 Guadalupe River, LL = Landa Lake, LSMR = Lower San Marcos River, and SMR_HW = San Marcos River headwaters.

## References

Abdelhady AA, Abdelrahman E, Elewa AM, Fan J, Zhang S, Xiao J (2018) Phenotypic plasticity of the gastropod *Melanoides tuberculata* in the Nile Delta: A pollution-induced stabilizing selection. Marine pollution bulletin 133:701–710

Albarrán-Mélzer NC, Ruiz LJR, Benítez HA, Lagos ME (2019) Can temperature shift morphological changes of invasive species? A morphometric approach on the shells of two tropical freshwater snail species. Hydrobiologia:1–10

Bandelt H, Forster P, Röhl A (1999) Median-joining networks for inferring intraspecific phylogenies. Mol Biol Evol 16:37–48

Chiu YW, Gan YC, Kuo PH, Hsu KC, Tan MS, Ju YM, Lin HD (2018) Mitochondrial genetic diversity of the freshwater snail Melanoides tuberculata. Biochemical Genetics:1–15

Coenye T, Vandamme P (2003) Intragenomic heterogeneity between multiple 16S ribosomal RNA operons in sequenced bacterial genomes. FEMS Microbiology Letters 228:45–49 doi:10.1016/s0378-1097(03)00717-1

Crowe J, Sharp Jr J (1997) Hydrogeologic delineation of habitats for endangered species: The Comal Springs/River system. Environmental Geology 30:17–28

Daniel W, Benson A, Neilson M (2019) Melanoides tuberculata (Muller, 1774): U.S. Geological Survey. Nonindigenous Aquatic Species Database, Gainesville, FL https://nas.er.usgs.gov/queries/FactSheet.aspx?SpeciesID=1037

Denver DR, Morris K, Lynch M, Vassilieva LL, Thomas WK (2000) High direct estimate of the mutation rate in the mitochondrial genome of Caenorhabditis elegans. Science 289:2342–2344

Effler SW et al. (1996) Impact of zebra mussel invasion on river water quality. Water Environment Research 68:205–214

Facon B, Jarne P, Pointier JP, David P (2005) Hybridization and invasiveness in the freshwater snail *Melanoides tuberculata*: Hybrid vigour is more important than increase in genetic variance. Journal of Evolutionary Biology 18:524–535

Facon B, MacHline E, Pointier JP, David P (2004) Variation in desiccation tolerance in freshwater snails and its consequences for invasion ability. Biological Invasions 6:283–293

Facon B, Pointier JP, Glaubrecht M, Poux C, Jarne P, David P (2003) A molecular phylogeography approach to biological invasions of the New World by parthenogenetic Thiarid snails. Molecular Ecology 12:3027–3039

Fleming PB, Huffman DG, Bonner TH, Brandt TM (2011) Metacercarial distribution of *Centrocestus formosanus* among fish hosts in the Guadalupe river drainage of Texas. Journal of Aquatic Animal Health 23:117–124

Freitas J, Bedê L, Júnior M, Rocha L, Santos M (1987) Population dynamics of aquatic snails in Pampulha reservoir. Memórias do Instituto Oswaldo Cruz 82:299–305

Genner MJ, Michel E, Erpenbeck D, De Voogd N, Witte F, Pointier JP (2004) Camouflaged invasion of Lake Malawi by an Oriental gastropod. Molecular Ecology 13:2135–2141

Gustafson K, Kensinger B, Bolek M, Luttbeg B (2014) Distinct snail *(Physa)* morphotypes from different habitats converge in shell shape and size under common garden conditions. Evolutionary Ecology Research 16:77–89

Haag-Liautard C, Coffey N, Houle D, Lynch M, Charlesworth B, Keightley PD (2008) Direct estimation of the mitochondrial DNA mutation rate in *Drosophila melanogaster*. PLoS biology 6:e204

Heller J, Farstey V (1990) Sexual and parthenogenetic populations of the freshwater snail *Melanoides tuberculata* in Israel. Israel Journal of Zoology 37:75–87 doi:10.1080/00212210.1990.10688643

Huelsenbeck JP, Ronquist F (2001) MRBAYES: Bayesian inference of phylogenetic trees. Bioinformatics 17:754–755

Karatayev AY, Burlakova LE, Karatayev VA, Padilla DK (2009) Introduction, distribution, spread, and impacts of exotic freshwater gastropods in Texas. Hydrobiologia 619:181–194

Keawjam R, Poonswad P, Upatham E, Banpavichit S (1993) Natural parasitic infection of the golden apple snail, *Pomacea canaliculata*. The Southeast Asian Journal of Tropical Medicine and Public Healt 24:170–177

Lindholm JT (1979) The gastropods of the upper San Marcos River and their trematode parasites. MS Thesis. Southwest Texas State University

Livshits G, Fishelson L (1983) Biology and reproduction of the freshwater snail *Melanoides tubercolata* (Gastropoda: Prosobranchia) in Israel. Israel Journal of Zoology 32:21–35

Livshits G, Fishelson L, Wise GS (1984) Genetic similarity and diversity of parthenogenetic and bisexual populations of the freshwater snail *Melanoides tuberculata* (Gastropoda: Prosobranchia). Biological Journal of the Linnean Society 23:41–54

Lynch M et al. (2008) A genome-wide view of the spectrum of spontaneous mutations in yeast. Proceedings of the National Academy of Sciences 105:9272–9277

Mitchell AJ, Brandt TM (2005) Temperature tolerance of red-rim melania *Melanoides tuberculatus*, an exotic aquatic snail established in the United States. Transactions of the American Fisheries Society 134:126–131

Mitchell AJ, Hobbs MS, Brandt TM (2007) The effect of chemical treatments on red-rim melania *Melanoides tuberculata*, an exotic aquatic snail that serves as a vector of trematodes to fish and other species in the USA. North American Journal of Fisheries Management 27:1287–1293

Murray H (1964) *Tarebia granifera* and *Melanoides tuberculata* in Texas. Annual Report to the American Malacological Union 53:15–16

Murray H (1971) The introduction and spread of Thiarids in the United States. The Biologist 53:133–135

Murray H (1975) *Melanoides tuberculata* (Müller), Las Moras Creek, Bracketville, Texas. Bulletin of the American Malacological Union 43

Murray H, Wopschall L (1965) Ecology of *Melanoides tuberculata* (Müller) and *Tarebia granifera* (Lamarck) in south Texas. Annual Reports 1965:25–26

Neck RW (1985) *Melanoides tuberculata* (Thiaridae) in extreme southern Texas. Texas Conchologist 21:150–152

Nelson NM (2019) Enumeration of potential economic costs of dreissenid mussels infestation in Montana. Universtiy of Montana http://dnrc.mt.gov/divisions/cardd/docs/misac-docs/dnrc_economic_cost_dreisseid_mussels_0119.pdf

Nollen P, Murray H (1978) *Philophthalmus gralli:* identification, growth characteristics, and treatment of an oriental eyefluke of birds introduced into the continental United States. The Journal of parasitology 64:178–180

Pinto HA, De Melo AL (2011) A checklist of trematodes (Platyhelminthes) transmitted by *Melanoides tuberculata* (Mollusca: Thiaridae). Zootaxa 28:15–28

Pointier J-P, Théron A, Borel G (1993) Ecology of the introduced snail *Melanoides tuberculata* (Gastropoda: Thiaridae) in relation to *Biomphalaria glabrata* in the marshy forest zone of Guadeloupe, French West Indies. Journal of Molluscan Studies 59:421–428

Pokora Z (2001) Role of gastropods in epidemiology of human parasitic diseases. Wiadomosci parazytologiczne 47:3–24

Rader RB, Belk MC, Keleher MJ (2003) The Introduction of an invasive snail *(Melanoides tuberculata)* to spring ecosystems of the Bonneville Basin, Utah. Journal of Freshwater Ecology 18:647–657

Ricciardi A, Neves RJ, Rasmussen JB (1998) Impending extinctions of North American freshwater mussels (Unionoida) following the zebra mussel *(Dreissena polymorpha)* invasion. Journal of animal ecology 67:613–619

Riley LA, Dybdahl MF, Hall Jr RO (2008) Invasive species impact: asymmetric interactions between invasive and endemic freshwater snails. Journal of the North American Benthological Society 27:509–520

Roessler M, Beardsley G, Tabb D (1977) New records of the introduced snail, *Melanoides tuberculata* (Mollusca: Thiaridae) in south Florida. Florida Scientist:87–94

Russo TN (1973) Discovery of the gastropod snail *Melanoides (Thiara) tuberculata* (Müller) in Florida. Florida Scientist 36:212–213

Sakai AK et al. (2001) The population biology of invasive species. Annual Review of Ecology and Systematics 32:305–332

Samadi S et al. (1998) Density and variability of dinucleotide microsatellites in the parthenogenetic polyploid snail *Melanoides tuberculata*. Molecular Ecology 7:1233–1236

Samadi S, David P, Jarne P (2000) Variation of shell shape in the clonal snail *Melanoides tuberculata* and its consequences for the interpretation of fossil series. Evolution 54:492–502

Samadi S, Mavárez J, Pointier JP, Delay B, Jarne P (1999) Microsatellite and morphological analysis of population structure in the parthenogenetic freshwater snail *Melanoides tuberculata:* Insights into the creation of clonal variability. Molecular Ecology 8:1141–1153

Saunders KS et al. (2001) An evaluation of spring flows to support the upper San Marcos River spring ecosystem, Hays County, Texas. Texas Parks and Wildlife Department. River Studies Report

Sørensen LVG, Jørgensen A, Kristensen TK (2005) Molecular diversity and phylogenetic relationships of the gastropod genus *Melanoides* in Lake Malawi. African Zoology 40:179–191

Thompson J, Higgins D, Gibson T (1994) CLUSTAL W: improving the sensitivity of progressive multiple sequence alignment through sequence weighting, position-specific gap penalties and weight matrix choice. Nucleic Acids Research 22:4673–4680

U.S. Congress Office of Technology Assessment (1993) Harmful non-indigenous species in the United States vol OTA-F-565. U.S. Government Printing Office, Washington, DC

Van Bocxlaer B, Clewing C, Etimosundja J-PM, Kankonda A, Ndeo OW, Albrecht C (2015) Recurrent camouflaged invasions and dispersal of an Asian freshwater gastropod in tropical Africa. BMC Evolutionary Biology 15:1–18

Wickham H (2016) ggplot2: Elegant graphics for data analysis. Springer-Verlag New York,

Wingard GL, Murray JB, Schill WB, Phillips EC (2008) Red-rimmed melania (Melanoides tuberculatus) - A snail in Biscayne National Park, Florida - harmful invader or just a nuisance? U.S. Geological Survey Fact Sheet 2008–3006. U.S. Geological Survey:1–6

Work K, Mills C (2013) Rapid population growth countered high mortality in a demographic study of the invasive snail, *Melanoides tuberculata* (Müller, 1774), in Florida. Aquatic Invasions 8:417–425

Xu S, Schaack S, Seyfert A, Choi E, Lynch M, Cristescu ME (2011) High mutation rates in the mitochondrial genomes of *Daphnia pulex*. Molecular Biology and Evolution 29:763–769

Yang L, Tan Z, Wang D, Xue L, Guan M-X, Huang T, Li R (2014) Species identification through mitochondrial rRNA genetic analysis. Sci Rep 4:4089–4089 doi:10.1038/srep04089

